# Integrative Inference of Spatially Resolved Cell Lineage Trees using LineageMap

**DOI:** 10.64898/2026.01.19.700383

**Authors:** Xinhai Pan, Yiru Chen, Xiuwei Zhang

**Affiliations:** School of Computational Science and Engineering, Georgia Institute of Technology, Atlanta GA 30332, USA

**Keywords:** Cell lineage inference, Spatial transcriptomics, Evolutionary modeling, Maximum likelihood, Spatio-temporal processes

## Abstract

Understanding the spatio-temporal processes of tissue growth, including how new cell types emerge and how cells form the tissue architecture, is a fundamental problem in biology. The emerging spatially resolved lineage tracing data, where three modalities, lineage barcodes, gene expression profiles, and spatial locations, are measured for each single cell, provides an unprecedented opportunity to understand these processes. Computational methods that take advantage of all three modalities to reconstruct cell lineage tree and ancestral cell states and locations are needed. We introduce LineageMap, a hybrid lineage inference algorithm that integrates the scalability of distance-based tree reconstruction methods with the flexibility of likelihood-based methods under a unified probabilistic framework. The input to LineageMap is spatially resolved lineage tracing data, where for each single cell, the gene expression, lineage barcode and spatial locations are available. LineageMap enables accurate, interpretable, and scalable inference of high-resolution lineage trees as well as locations of ancestral cells from the tri-modality single-cell data. Across simulated and experimental datasets, LineageMap consistently outperforms existing methods in the accuracy of reconstructed cell lineage trees, while revealing biologically coherent spatiotemporal trajectories. Our framework bridges molecular lineage tracing with spatial and transcriptomic information, advancing computational reconstruction of dynamic cellular ancestries in both time and space. LineageMap is available at: https://github.com/ZhangLabGT/LineageMap.

## 1 Introduction

Understanding how cells divide, differentiate, and organize into tissues and organs is a central question in developmental biology. Lineage tracing technologies aim to reconstruct the *cell division history*, or lineage tree, of a tissue by recording heritable molecular marks that accumulate during successive cell divisions. Recent advances in sequencing-based lineage tracing,^23, 28^ particularly those leveraging CRISPR/Cas9 genome editing, have enabled joint measurement of lineage barcodes and transcriptomes across thousands of single cells.^29^ These barcodes consist of engineered genomic sequences—lineage recorders—that undergo stochastic mutations at designated target sites over successive cell divisions,^1, 4, 25^ that can be used to reconstruct detailed lineage trees to uncover latent developmental trajectories and division hierarchies that cannot be directly observed *in vivo*. In a standard lineage tree, each node represents a single cell, and each edge denotes a direct parent–daughter division event.

More recently, a new wave of spatiotemporal lineage tracing datasets are emerging,^12–14, 30^ which integrate lineage tracing with spatial transcriptomics technologies. This combination enables the simultaneous observation of cell lineage barcodes, expression and spatial locations at single-cell resolution, which respectively represent cell division, differentiation, and spatial organization. Such tri-modality datasets make it possible to reconstruct not only the ancestral relationships among cells, but also the spatial trajectories of cell divisions and migrations – giving rise to the concept of *spatially resolved lineage trees*. These results offer unprecedented opportunities to understand the coordination between lineage, spatial architecture and cell fate.

However, reconstructing spatially resolved lineage trees poses substantial computational challenges. Existing lineage reconstruction methods focus primarily on optimizing the topology of cell division trees from barcode mutations,^9, 22^ and some recent methods use barcode mutations and single cell gene expression data.^18, 31^ Without considering spatial information of cells, these methods do not take advantage of the mutual constraints between lineage, spatial proximity, and cell state continuity — key factors in developmental processes. To fully leverage these new datasets, there is a need for integrative frameworks that jointly infer lineage structure while modeling spatial coherence and state transitions.

Classical Neighbor Joining (NJ)^21^ efficiently infers lineage tree topology based on pairwise distances between lineage barcodes. However, it assumes all taxa (cells) are distinct and that distances faithfully capture ancestral relationships. In real lineage barcode data, this assumption is violated due to dropout events and sparse mutation patterns, where many loci remain unedited or ambiguously sequenced, while *homoplasy*,^27^ where identical or near-identical barcodes, arise independently in different cells. On the other hand, maximum-likelihood (ML)^7^ or Bayesian tree inference methods^10, 32^ are becoming computationally intractable for the ever-increasing number of cells sequenced in lineage tracing experiments, as the number of possible unrooted trees increases in the order of *n*!! for *n* taxa. Furthermore, local search algorithms used in likelihood optimization are prone to convergence toward local optima, particularly in high-dimensional barcode spaces.^8, 15, 16, 26^ These challenges render full ML inference impractical for modern lineage recording datasets comprising thousands of cells.

Besides these challenges, the *polytomy* problem^19, 24^ is also critical, where many cells share identical barcodes causing one internal node having degree greater than three in an unrooted tree, additional information sources such as spatial proximity or transcriptomic similarity can be integrated to enhance the distinguishability of cells and to refine local tree structures.

To this end, taking advantage of the cutting-edge spatially-resolved lineage tracing datasets, we developed LineageMap, a hierarchical hybrid framework that integrates the computational scalability of distance-based inference with the accuracy and flexibility of likelihood-based optimization. By coupling statistical modeling with multi-modal context, LineageMap enables accurate and interpretable reconstruction of high-resolution lineage trees from large-scale, dropout-prone single-cell datasets, paving the way for deeper insights into the spatial and temporal dynamics of development, regeneration, and disease progression. Our tests on both simulated and real datasets demonstrated that LineageMap outperforms state-of-the-art methods for lineage tree reconstruction. It is worth mentioning that LineageMap differs fundamentally from transcriptomics-only trajectory inference methods such as RNA velocity or pseudotime inference methods. These approaches aim to infer transcriptional progression rather than true clonal ancestry. Because LineageMap uses gene expression as a prior to guide tree topology consistent with barcode mutations, benchmarking against trajectory inference methods would conflate distinct objectives and introduce circularity. Therefore, we will only focus on comparisons of lineage reconstruction methods in this paper.

## 2 Methods

### 2.1 Overview of LineageMap

In order to optmize computational cost and enhance robustness against technical noise in the barcode data, LineageMap first aggregates cells with highly similar lineage barcodes by applying Louvain clustering on a similarity graph derived from a dropout-aware weighted Hamming distance matrix on the barcodes, then obtains a cluster-level consensus barcode for each cluster, and applies Neighbor Joining to the consensus barcodes, which yields a backbone tree of cell clusters (Fig. 1).

**Fig. 1.**
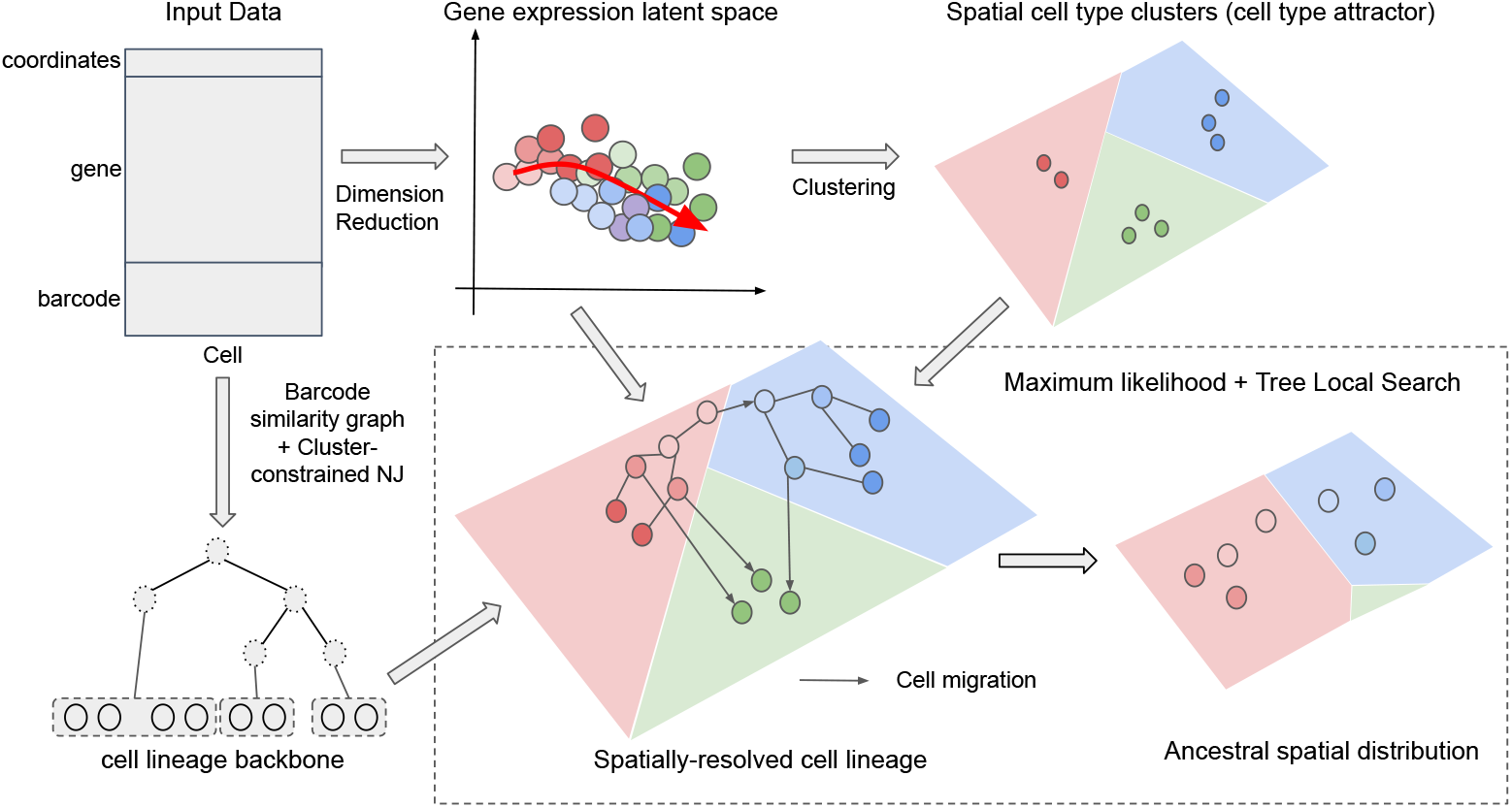
Overview of LineageMap algorithm. LineageMap integrates spatial coordinates, gene expression, and DNA barcodes to infer spatially resolved cell lineages. Gene expression is embedded and clustered to define cell-state groups, while a barcode-similarity graph with cluster-constrained Neighbor Joining provides an initial tree backbone. Using a maximum-likelihood framework with local tree search, LineageMap refines this backbone by incorporating spatial constraints and cluster structure to produce a spatially resolved lineage and its inferred ancestral spatial distribution.

LineageMap then performs maximum likelihood optimization within each cluster to determine the tree topology and branch lengths for these cells using multi-modal information, including barcode states, spatial coordinates, and cell states (Fig. 1). By modeling processes of barcode mutations, cell type differentiation and cells’ spatial localization during cell divisions, the likelihood function couples these relevant constraints, thus the three modalities together are expected to lead to more accurate reconstructed cell lineage trees with higher resolution, compared to methods using only the barcode or the barcode and gene expression data. In principle, the maximum likelihood framework can be used to reconstruct the whole lineage tree without building the backbone first. However, for lineage tracing data with thousands of cells, the bottleneck is not likelihood evaluation itself but the combinatorial tree topology search. Existing barcode-based ML approaches such as LinTIMaT,^31^ LinRace,^18^ and BiLinT,^5^ also tend to adopt heuristic or constrained search strategies to reduce the effective search space.

In LineageMap, the initial distance-based backbone clustering serves to partition cells into coarse clonal groups that share highly similar barcodes, thereby reducing topological ambiguity caused by early barcode noise. The subsequent probabilistic refinement is then performed within each cluster, where the search space is dramatically reduced and likelihood-based local search becomes more efficient. Empirically, we observed that direct global ML search without this backbone initialization frequently becomes trapped in poor local optima and exhibits substantially higher runtime. Thus, the two-stage design improves both computational efficiency and stability of inference.

### 2.2 Cell lineage tree

We define a rooted, directed, bifurcating cell lineage tree *T* = (*V, E*) with leaf set corresponding to the observed cells, internal nodes representing unobserved ancestors, and branch lengths { *t*_*uv*_ } _(*u*→*v*)∈*E*_ proportional to elapsed time. An edge (*u* → *v*) represents a parent–daughter cell relationship. Let **b**_*v*_, **s**_*v*_ and **z**_*v*_ denote the (latent) barcode, spatial coordinates, and cell state of node *v*.

### 2.3 Spatially-resolved lineage tree inference problem

Let *n* be the number of observed single cells. For each cell *i* ∈ {1, …, *n*} we observe:

– a lineage barcode **b**_*i*_ = (*b*_*i*1_, …, *b*_*iL*_) across *L* target sites,
– a spatial coordinate **s**_*i*_ ∈ ℝ^*d*^,
– a categorical cell state label **z**_*i*_ ∈ {1, …, *C*}.

We seek a rooted tree *T* = (*V, E*) that best explains the observed tri-modality data through spatio-temporal cell division and localization processes. We first detect lineage clones and construct a backbone tree. We then formulate generative models for lineage barcodes, spatial locations, and cell states, and derive a joint likelihood function (Eq. (3)) under a Bayesian framework.

### 2.4 Detection of lineage clones and tree backbone inference

To address the lineage polytomy problem, we first identify coherent *lineage clones*—groups of cells sharing highly similar or identical barcodes—using a community detection approach on the barcode similarity graph. Specifically, we apply Louvain clustering with a tunable resolution threshold to partition cells into candidate clonal groups. Each clone represents a local subtree that can be independently refined before integrating into a global lineage topology. The threshold parameter should be tuned to the dropout level in the data, and the detailed description of determining the threshold value can be found in Supplementary Material.

We then infer a coarse-grained *backbone tree* representing the relationships among clones. We adapt the standard NJ method to reconstruct this tree while ensuring cells in the same clone to form a subtree. We name this NJ method the “cluster-constrained NJ” method. The resulting tree is a backbone tree that provides the overall topology while allowing fine-grained optimization within the clone.

Finally, each clone subtree is inferred in parallel using a stochastic optimization framework that maximizes a likelihood function, that jointly models the evolution of barcode, cell states and cell locations. After local refinement, all subtrees are merged back into the backbone tree by replacing clone tips on the backbone with their corresponding optimized subtrees.

### 2.5 Generative models of spatio-temporal lineage trees Barcode evolution model

We model the evolution of CRISPR-Cas9 lineage barcodes as an irreversible mutational continuous-time Markov chain (CTMC) process on a finite set of editing outcomes. Each barcode consists of *L* independent target sites (“cut sites”), indexed by *l* = 1, …, *L*. For cell *i*, the observed barcode is **b**_*i*_ = (*b*_*i*1_, …, *b*_*iL*_), where *b*_*il*_ ∈ 𝒜_*l*_, and 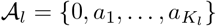 denotes the character space of locus *l*. Here, 0 corresponds to the unedited state, while 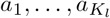 correspond to distinct irreparable mutations (“scars”) generated by Cas9-induced indels. Additionally, an extra symbol ∅ is included to represent dropout or missing observations.

The underlying CTMC starts in state 0 at the root cell. A character in the 0 state of the parent barcode mutates at a rate of 0 *< λ <* 1. When a character in the 0 state mutates, it acquires a new state from the set 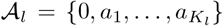, where the state *a*_*i*_ is acquired with probability *P* (*a*_*i*_). This way, the CTMC is parameterized by *λ >* 0 and the state probabilities 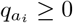. We denote 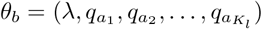.

#### Spatial evolutionary model of cell division and differentiation

For a daughter cell *v* that divided from a parent cell *u*, the spatial location of *v, s*_*v*_, follows a diffusion model along the tree,

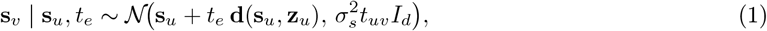

where **d**( · ) is a possible drift field (e.g., movement along tissue axes or chemotactic gradients), and 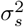 is spatial diffusivity. We allow **d** to depend on the cell state **z**_*u*_, which means cells in different states migrate differently. *t*_*uv*_ is the edge length of edge (*u, v*). *I*_*d*_ is an *d × d* identity matrix, where *d* = 2.

#### Brownian motion model of cell states

We model the cell state of node *v*, **z**_**v**_, as a multivariate trait evolving via a Brownian motion process, depending on the cell state of its parent, **z**_**u**_:

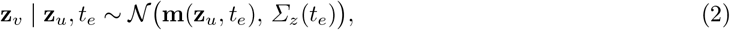

where **m**(**z**_*u*_, *t*_*uv*_) = **z**_*u*_ + *t*_*e*_ *β*(**z**_*u*_) can include drift toward attractor states (lineage-specific differentiation directions).

### 2.6 Joint likelihood and posterior

The observed data is 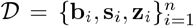, which represent the lineage barcodes, spatial locations and cell states of leaf cells. The joint likelihood given (*T*, {*t*_*e*_}, *θ*) factorizes, with latent node variables marginalized:

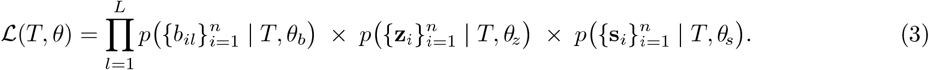

The three terms that multiply are respectively the likelihood of lineage barcodes, cell states and cell locations on a lineage tree *T* . These three terms are detailed in following sections. The posterior probability then becomes:

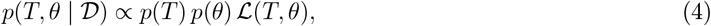

where *p*(*T* ) is the prior for the tree candidate *T* and *p*(*θ*) includes priors for the latent parameters, including barcode mutation rate, state probabilities, and spatial drift parameters.

### 2.7 Barcode likelihood

Let *T* = (*V, E*) be a rooted tree with root *r*, internal nodes *V*_int_, and leaves corresponding to the *n* observed cells. Each leaf *i* has an observed barcode **b**_*i*_ = (*b*_*i*1_, …, *b*_*iL*_) across *L* independent loci. For locus *l*, the finite state space is 𝒜_*l*_ as defined earlier. Denote by *t*_*uv*_ the length of edge (*u* → *v*) ∈ *E*.

#### Barcode state transition model

For locus *l*, edits occur irreversibly from the unedited state. That is, once a locus is edited, its state is locked and will not mutate again. Let *λ*_*lk*_ be the rate of editing from un-mutated state 0 to a mutated state *k*, with total mutation rate 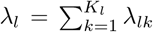. On an edge of length *t*, following Camin-Sokal Parsimony,^2^ we can derive the transition probabilities between barcode states: Thus, for edge (*u* → *v*) of length *t*_*uv*_,

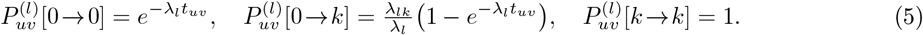

A matrix form of the transition probabilities is included in Supplementary Material.

Given the transition probabilities, we use a bottom-up approach to recursively calculate the barcode likelihood on the whole tree. Define the likelihood of a subtree rooted at node *v* at locus *l* as 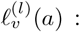 = Pr(observations in subtree rooted at *v* | *v* has character *a*) .

If *v* is a leaf node, denote the observation by *o*_*vl*_ ∈ 𝒜_*l*_ ∪{∅}, where ∅ denotes dropout. In case of dropout events in the data, 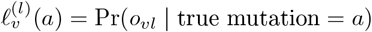; If there is no dropout, 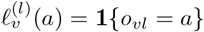.

If *v* is an internal node with children ch(*v*), we have:

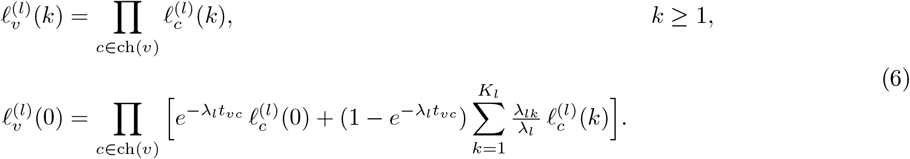

When we get to the root *r*, the likelihood 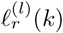 becomes the likelihood of the whole tree for locus *l*. With prior 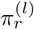 on the root, the likelihood on locus *l* can be calculated as 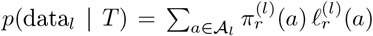 If the root is fixed to the unedited state, such as in the CRISPR/Cas9 model, the per-locus likelihood is 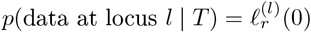.

Assuming loci evolve independently, the total likelihood is

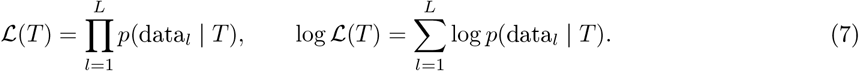

### 2.8 Likelihood of state and space evolution

We next consider the joint evolution of discrete cell states and continuous spatial locations of cells that evolve following *T* = (*V, E*) with root *r* and edge lengths {*t*_*uv*_}. Leaves *i* = 1, …, *n* have observed cell state *z*_*i*_ and continuous spatial coordinate *s*_*i*_ ∈ ℝ^*d*^. The joint likelihood of cell states and locations of leaf cells is

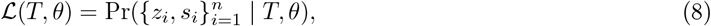

computed by marginalizing over unobserved internal node variables. When deriving this likelihood, we use CTMC to model cell state evolution and Gaussian message passing to model cells’ spatial location evolution.

#### Discrete state evolution using CTMC

Cell states *z* ∈ { 1, …, *C* } evolve along the tree as a continuous-time Markov chain with rate matrix *R*. For edge (*u* → *v*) of length *t*_*uv*_, we define the transition probability matrix *M*, where

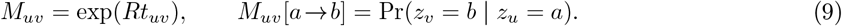

We then use a bottom-up recursive procedure to calculate the cell state likelihood on the tree, similar to what we do with the barcode likelihood. The recursion aggregates subtree likelihoods by summing over child state transitions and multiplying across independent branches. We define *ℓ*_*v*_(*b*), as the likelihood of a subtree rooted at node *v*. When *v* is a leaf, we have *ℓ*_*v*_(*b*) = *e*_*v*_(*b*), where *e*_*v*_(*b*) = Pr(*o*_*v*_ | *z*_*v*_ = *b*) if there may be dropouts, and *ℓ*_*v*_(*b*) = **1**{*o*_*v*_ = *b*} in dropout-free data.

Then for internal node *v* with state *a* and children ch(*v*), we have:

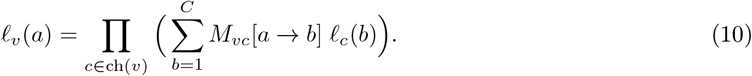

Finally, for root node *i* with root prior *π*_*r*_, we have

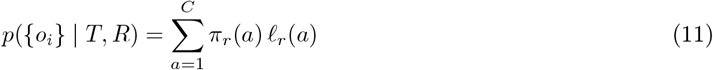

#### Spatial coordinate evolution with Gaussian message passing

Let *s* ∈ ℝ^*d*^ denote continuous spatial coordinates. Along an edge (*u* → *v*) of length *t*, we use a Brownian motion model for the shift of daughter cells from the parent cell:

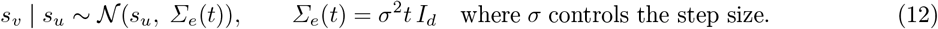

#### Message passing

Suppose child *c* contributes message *p*(data in subtree *c* | *s*_*c*_) ∝ 𝒩 (*s*_*c*_; *m*_*c*_, *S*_*c*_). Integrating over *s*_*c*_ yields a Gaussian in *s*_*v*_, we have:

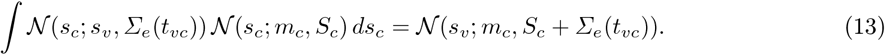

Thus each child contributes a Gaussian distribution in *s*_*v*_, and their product is still a Gaussian distribution. *Information form update*. For child *c*, define

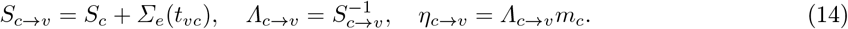

Aggregating across children, we have:

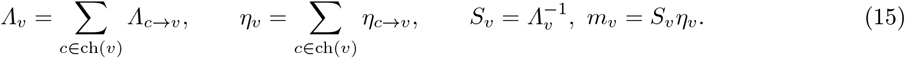

#### Likelihood at the root

If the root has prior *s*_*r*_ ∼ 𝒩 (*µ*_0_, Σ_0_), then with message 𝒩 (*s*_*r*_; *m*_*r*_, *S*_*r*_) from the subtree,

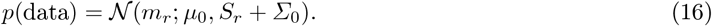

### 2.9 State-dependent spatial OU model with lineage-specific attractors

In order to model cell migration, we extend the Brownian motion model to Ornstein–Uhlenbeck(OU) model (Details about the extension derivations can be found in Supplementary Material). Let *T* = (*V, E*) be a rooted tree with branch (*u* → *v*) of length *t*_*uv*_ *>* 0. Each node *v* has a *d*-dimensional spatial coordinates **s**_*v*_ ∈ ℝ^*d*^ and a cell state *z*_*v*_ ∈ { 1, …, 𝒞 }(observed at leaves, latent at internals). Dynamics along an edge are governed by the *parent* state *z*_*u*_.

For each state *z* ∈ {1, …, 𝒞}, we posit a scalar OU rate *α*_*s*_ *>* 0, a scalar diffusion 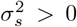 (isotropic covariance), and an attractor ***µ***_*s*_ ∈ ℝ^*d*^. The OU transition on edge (*u* → *v*) is

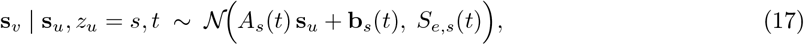

with

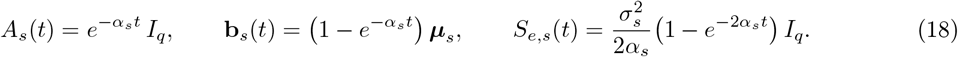

For *t* = 0 use the continuous limits *A*_*s*_(0) = *I*_*q*_, **b**_*s*_(0) = **0**, *S*_*e,s*_(0) = 0.

#### Upward message from child c to parent u

Suppose the child’s upward message is Gaussian 𝒩 (**m**_*c*_, *S*_*c*_). Let *W*_*s*_ (*t*) ≡ (*S*_*c*_ + *S*_*e,s*_ (*t*))^−1^. With *A* = *A*_*s*_ (*t*) and **b** = **b**_*s*_ (*t*), the child contribution on **s**_*u*_ is Gaussian with information parameters

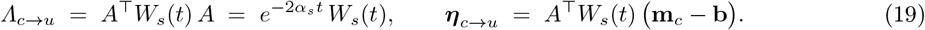

At a parent *u* with children ch(*u*), combine by summation

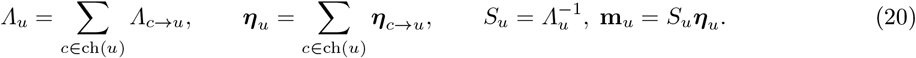

At the root *r*, combine with a Gaussian prior 𝒩 (***µ***_0_, Σ_0_) in the usual way:

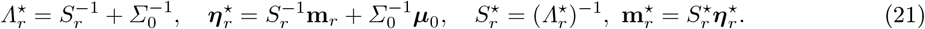

#### Marginal spatial log-likelihood (accumulated)

During the upward pass, add

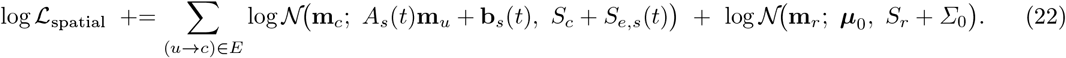

The posterior means {**m**_*u*_}_*u*∈*V*_ provide the inferred internal coordinates.

### 2.10 Construct the full lineage tree and whole tree refinement

The joint likelihood in Eq. 3 is used to optimize subtrees for each cell clone, using a local search algorithm. We adopted random Subtree Swapping (rSS), which is a derivative of the widely used random Subtree Pruning and Regrafting (rSPR).^6^ We randomly select two nodes on the current tree and prune the subtrees attached to the specific nodes. Then, we regraft either one of the pruned subtrees to the location of the other subtree. Then, we adopt a first-improvement local search strategy using rSS. While subtree prune-and-regraft (SPR) is a standard phylogenetic move, exhaustive evaluation of all SPR neighbors becomes computationally expensive for large clones. Instead, we sample candidate subtree swaps and accept the first likelihood-improving move encountered. This stochastic first-improvement strategy substantially reduces runtime while maintaining effective exploration of the topology space.^18^

With the inferred tree backbone and the optimized subtrees, LineageMap builds the initial full lineage by re-attaching the subtrees onto the tree backbone at the leaves. In the end, we run an additional whole tree likelihood evaluation to infer ancestral state and spatial coordinates.

### 2.11 Simulating synthetic datasets with SpaTedSim

To quantitatively evaluate LineageMap and other methods under controlled and biologically realistic conditions, we developed SpaTedSim (Spatio-Temporal dynamics Simulation of single cells), a simulator that jointly models cell division, cell state shifts, and cellular movement in 2-D space (details in Supplementary Material and Supplementary Fig. 1). SpaTedSim directly simulates how the cell population grows from a single cell to observed cells with various cell types. With the modeling of the cell state tree, cell lineage tree, and spatial movements of cells, SpaTedSim generates realistic, paired gene expression, lineage barcodes, and spatial coordinates, and with the ground truth, we can look into the spatial distribution of cell types and lineages (Supplementary Fig. 1b) and uncover the spatio-temporal dynamics of cells, for example, examine spatial distribution of cells along the cell lineage tree (Supplementary Fig. 1c-d). SpaTedSim consists of two major steps: 1. Simulating cells’ lineage barcode and gene expressions on the cell division tree; 2. Simulating cells’ spatial coordinates based on the cell division tree.

## 3 Results

### 3.1 LineageMap outperforms existing methods on synthetic datasets

SpaTedSim provides complete ground-truth lineage trees and spatial coordinates for every cell, enabling quantitative assessment of reconstruction accuracy. Given the observed cells’ spatial locations (Fig. 2a), LineageMap can infer spatially-resolved lineage tree and infer ancestral spatial locations (Fig. 2b). We can compare the inferred lineage tree with the ground truth and benchmark against other methods.

**Fig. 2.**
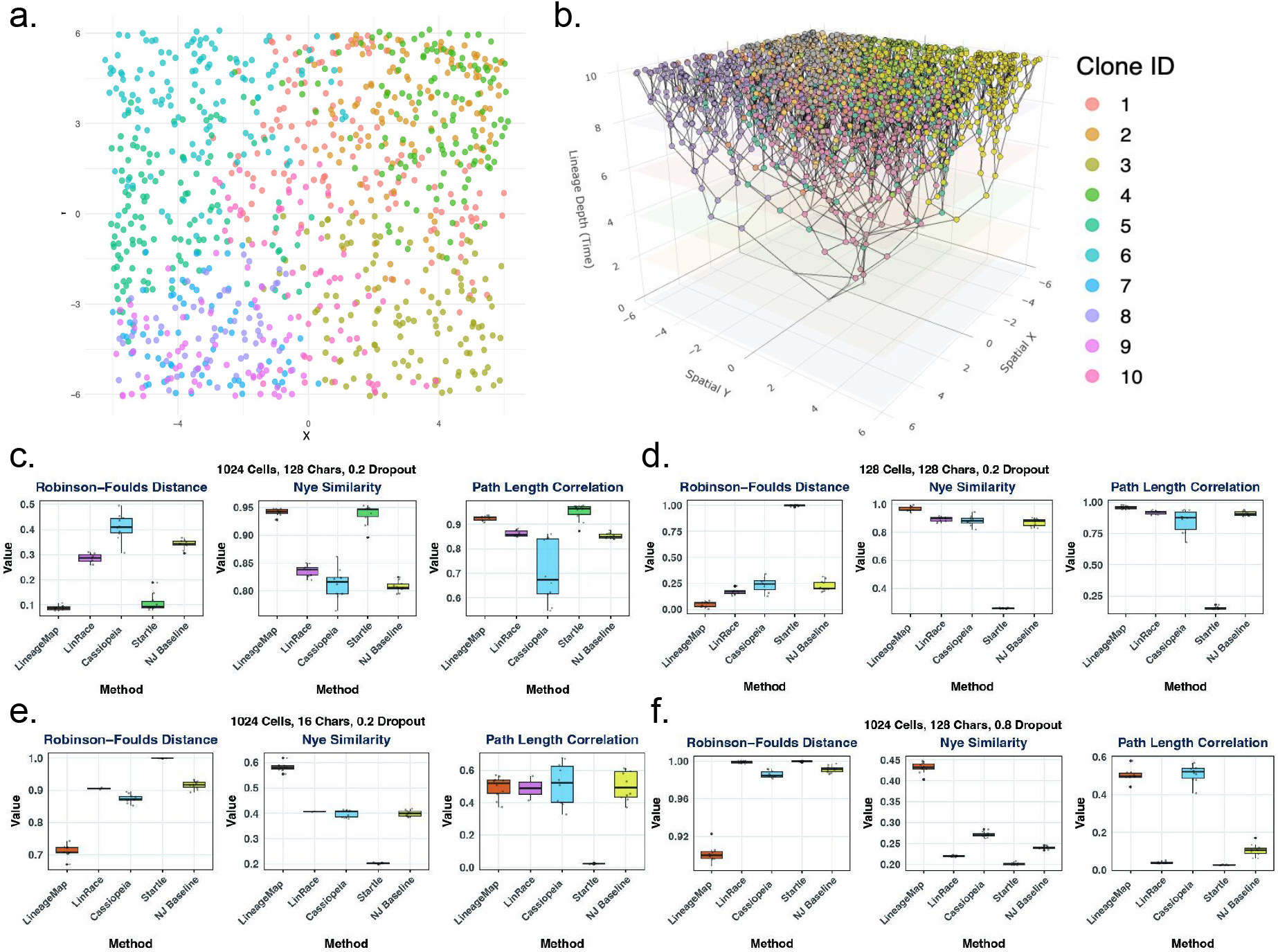
(a–b) Visualization of the default simulation dataset (replicate 1).(a) 2D scatter plot of cell spatial coordinates (X–Y), colored by Clone ID (major sub-lineages from hierarchical clustering on the reconstructed tree; average linkage on cophenetic distances, k=10), showing clear spatial segregation.(b) Plotly 3D visualization of the reconstructed lineage tree, with X–Y denoting spatial position, Z denoting evolutionary depth, and colors indicating clone ID.(c–f) Benchmark results for LineageMap, NJ, Cassiopeia (two modes), and Startle across simulated conditions. LineageMap consistently achieves the best or near-best performance in RF distance (lower is better), Nye similarity, and pathlength correlation (higher is better). Panels show results for (c) 1024 cells, 128 characters, 0.2 dropout; (d) 128 cells, 128 characters, 0.2 dropout; (e) 1024 cells, 16 characters, 0.2 dropout; and (f) 1024 cells, 128 characters, 0.8 dropout. Under high dropout (f), LineageMap shows markedly superior robustness.

We benchmarked LineageMap against state-of-the-art methods: LinRace,^18^ which uses both lineage barcodes and gene expression; Cassiopeia,^11^ a parsimony-based heuristic; Startle,^22^ a technique based on star homoplasy evolutionary model, and an NJ baseline on the weighted Hamming distances. We tested the methods across multiple simulation regimes that vary in three hyperparameters: the number of cells, target sites, and dropout rates (Fig. 2c-f). These conditions capture the major factors of difficulty in barcode-based lineage reconstruction. Evaluation metrics include Robinson–Foulds (RF) distance,^20^ Nye similarity,^17^ and path-length correlation, capturing complementary aspects of tree topology and branch-length structure (details on the metrics are in Supplementary Material).

In the benchmark setting with 1024 cells, 128 characters and 20% dropouts (Fig. 2c), LineageMap achieves the best overall performance, having the best RF distance and Nye Similarity, and second to Startle for path length correlation. To comprehensively compare the performances of the methods under various conditions, in Fig. 2d-f, we further compare the methods with a smaller number of cells (128 cells), a smaller number of target sites (16 target sites), or higher dropout rate (up to 80%).

When the number of cells is reduced, most methods improve due to the reduction in tree complexity. Nevertheless, LineageMap maintains leading performance across all three metrics, with LinRace being the second best. Surprisingly, Startle fails to recover meaningful topology, producing near-random topologies revealed by RF distance and Nye Similarity.

With only 16 target sites, the mutation signal becomes extremely sparse, creating a challenging inference scenario. As expected, all methods show reduced accuracy, but the performance gap widens. LineageMap again achieves the best RF and Nye scores, recovering substantially more topological structure than any baseline. For path length correlation, LinRace, Cassiopeia and NJ remains relatively similar performances as LineageMap. These results highlight that LineageMap is uniquely effective in low-information settings, preserving stable reconstruction accuracy even when mutations are scarce.

With a high dropout rate as 80%, most mutations are masked which creates another complex inference scenario. LineageMap remains the top performer by a large margin, with the lowest RF distance, highest Nye similarity and the best path length correlation alongside Cassiopeia. However, Cassiopeia, alongside with other methods, fall short in terms of inferring the overall topology. One thing to mention is that we observe a severe performance collapse of Startle from the default settings to the varied settings, and we think that such inconsistency might be due to some fatal error in the program or the model’s unable to handle different barcode qualities. We also performed additional analysis on varying dropout rates (0, 0.2, 0.4, 0.6, 0.8) to characterize the trend of the performances when the barcode data gets increasingly corrupted (Supplementary Fig. 2). Across three metrics, LineageMap is the most consistent and best performing method, especially when the dropout rate are higher, presenting the best robustness ( ≥ 0.4).

Across all four benchmarking conditions—varying cell count, barcode complexity, and dropout - LineageMap consistently achieves the best or near-best performance on every metric, with notably higher stability across replicates. The method performs particularly strongly in the most challenging scenarios (low target-site count and high dropout), where competing tree-reconstruction algorithms frequently fail to recover meaningful structure. We note that in simulation settings with extremely weak spatial-lineage coherence, the spatial prior may introduce noise rather than signal, leading to performance comparable to barcode-only methods. This behavior highlights that LineageMap does not artificially over-weight spatial information. In the end, these results show that LineageMap is a robust and accurate approach for lineage reconstruction across a wide range of experimental settings.

### 3.2 LineageMap reconstructs spatially-resolved lineage tree of baseMEMOIR mESC cell population

baseMEMOIR^3^ is a method that combines CRISPR base editing with in situ imaging of engineered “recording” arrays and sequential hybridization readout, allowing high-resolution recording of lineage barcodes in spatial context. Because baseMEMOIR provides spatially-indexed barcode data, it offers a direct testbed for benchmarking the ability of our algorithm (LineageMap) to resolve complex lineages in space, to test polytomy resolution, and to validate spatial-state coupling in inferred trees.

Given the observed lineage barcode, cell state, and spatial location of single-cells on grown mESC dishes, we use LineageMap to infer the lineage tree and the ancestral spatial locations at the same time (Fig. 3a). This spatially resolved lineage tree can illustrate how the population is grown from a single mESC cell at the root to the observed leaf cells. If we compress all generations into a 2D visualization (Fig. 2b-c), we can see that LineageMap can preserve the spatial clustering of each cell type, with naive mESC cells in the middle and formative mESC cells around the boundary of the population. Results on other samples can be found in the Supplementary Fig. 3-4.

**Fig. 3.**
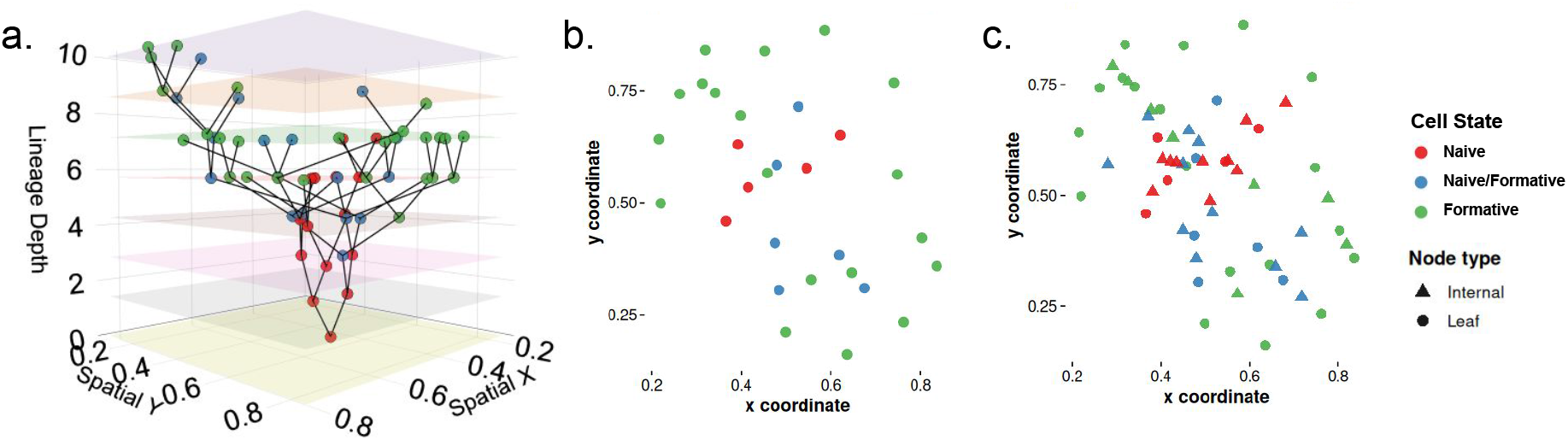
LineageMap infers spatially-resolved lineage on baseMEMOIR mESC data (colony 2). (a) A ploty 3D visualization of the reconstructed lineage tree projected into spatial and temporal dimensions. X and Y axes represent the spatial coordinates of the cells, while Z axis represents the evolutionary depth from the root, and cells are colored with cell types. (b) 2D visualization of spatial coordinates of observed leaf cells, colored by cell states. (c) 2D visualization of spatial coordinates of both observed leaf cells, and inferred ancestral cells, colored by cell states. The cell states and spatial coordinates of ancestral cells are inferred using LineageMap’s likelihood function.

We also performed a quantitative comparison between the inferred lineage trees and the tree reported in the original study. It serves as a reference tree that presents the phylogenetic topology inferred from both lineage barcodes and spatial locations. As shown in Table 1, LineageMap exhibits the highest agreement with the reference tree across the three similarity metrics. This observation suggests that incorporating spatial and gene expression information can help reduce ambiguities in branching structure and produce lineage trees that are more consistent with the spatial pattern, obtained as consensus from multiple trees and reflecting cell type spatial distribution, as shown in the baseMEMOIR paper.^3^

**Table 1.**
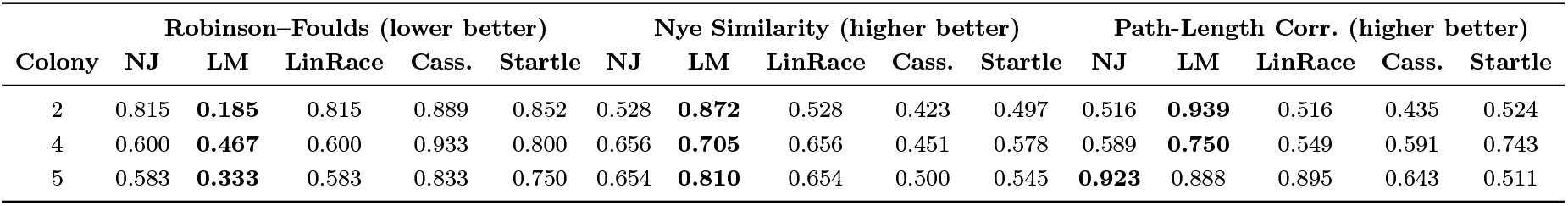
Comparing reconstructed trees across colonies for all methods and metrics.

| Colony | Robinson–Foulds (lower better) |  |  |  |  | Nye Similarity (higher better) |  |  |  |  | Path-Length Corr. (higher better) |  |  |  |  |
| --- | --- | --- | --- | --- | --- | --- | --- | --- | --- | --- | --- | --- | --- | --- | --- |
|  | NJ | LM | LinRace | Cass. | Startle | NJ | LM | LinRace | Cass. | Startle | NJ | LM | LinRace | Cass. | Startle |
| 2 | 0.815 | **0.185** | 0.815 | 0.889 | 0.852 | 0.528 | **0.872** | 0.528 | 0.423 | 0.497 | 0.516 | **0.939** | 0.516 | 0.435 | 0.524 |
| 4 | 0.600 | **0.467** | 0.600 | 0.933 | 0.800 | 0.656 | **0.705** | 0.656 | 0.451 | 0.578 | 0.589 | **0.750** | 0.549 | 0.591 | 0.743 |
| 5 | 0.583 | **0.333** | 0.583 | 0.833 | 0.750 | 0.654 | **0.810** | 0.654 | 0.500 | 0.545 | **0.923** | 0.888 | 0.895 | 0.643 | 0.511 |

## 4 Discussion

In this paper, we developed a new computational method, LineageMap, to infer spatially-resolved lineage trees from jointly profiled lineage barcode, gene expression and spatial coordinates data. This design specifically addresses core challenges in lineage reconstruction, including utlizing spatial location modality, high dropout rates, sparse mutation profiles, homoplasy, and barcode-induced polytomies.

LineageMap introduces two main advances: (i) barcode-based community detection combined with a cluster-constrained Neighbor Joining strategy to generate an efficient and robust global topology, and (ii) a unified likelihood model that integrates irreversible barcode mutations, spatial evolution, and cell-state transitions to refine local subtrees and infer ancestral states and locations.

Results on synthetic datasets demonstrate that LineageMap achieves consistent improvements across all major evaluation metrics, particularly in regimes where barcode quality decreases due to factors such as dropouts. Competing methods often show degraded or unstable performance under these challenging conditions, whereas LineageMap maintains both accuracy and robustness—indicating that spatial and transcriptomic information provide meaningful complementary signal for resolving ambiguous lineage relationships.

Analysis of the baseMEMOIR dataset further shows the utility of LineageMap in real experimental settings. The method reconstructs lineage topologies that better agree with consensus phylogenies while inferring spatially coherent ancestral positions and state transitions. This joint inference reveals the correspondence between lineage structure and spatial organization, a property that is inaccessible to barcode-only methods.

Current limitations include simplified spatial models that may not fully capture tissue-specific migration patterns. Future extensions incorporating richer spatial priors, accelerated likelihood evaluation, or continuous latent-state models may expand applicability and scalability. Additionally, current lineage tracing technologies typically profile several thousand cells per experiment. While emerging methods are increasing throughput, lineage barcode recovery efficiency remains a limiting factor. With more than 10K cells present, lineageMap can scale better than other state-of-the-art methods with the joint tree backbone + local refinement approach to reduce the sample size of each step.

Overall, LineageMap provides a scalable and statistically principled solution for reconstructing spatially resolved lineage trees from emerging tri-modality single-cell datasets. Although publicly available spatially resolved lineage tracing datasets are currently not widely available, given the increasing number of publications on such data,^12–14, 30^ methods that integrate lineage, spatial, and cellular-state information will play an essential role in high-resolution analysis of developmental and tissue dynamics in the coming future.

## 5 DATA AVAILABILITY

LineageMap is available at: https://github.com/ZhangLabGT/LineageMap.

SpaTedSim is a simulator for paired lineage barcode,spatial location and gene expression data available at: https://github.com/ZhangLabGT/SpaTedSim.

Cassiopeia is an end-to-end pipeline for single-cell lineage tracing experiments available at: https://github.com/YosefLab/Cassiopeia.

Startle is a set methods for lineage tree reconstruction by infering the most parsimonious tree under the star homoplasy evolutionary model available at: https://github.com/raphael-group/startle.

LinRace is a R package for lineage tree reconstruction, which can integrate the lineage barcode and gene expression data available at: https://github.com/ZhangLabGT/LinRace

## 6 FUNDING AND ACKNOWLEDGMENT

This work was supported by National Institutes of Health grant R35GM143070.

## References

1. Alemany, A., Florescu, M., Baron, C.S., Peterson-Maduro, J., van Oudenaarden, A.: Whole-organism clone tracing using single-cell sequencing. Nature 556(7699), 108–112 (Apr 2018). 10.1038/nature25969, https://doi.org/10.1038/nature25969

2. Camin, J.H., Sokal, R.R.: A method for deducing branching sequences in phylogeny. Evolution 19(3), 311–326 (Sep 1965). 10.1111/j.1558-5646.1965.tb01722.x, http://dx.doi.org/10.1111/j.1558-5646.1965.tb01722. x

3. Chadly, D.M., Frieda, K.L., Gui, C., Klock, L., Tran, M., Sui, M.Y., Takei, Y., Bouckaert, R., Lois, C., Cai, L., Elowitz, M.B.: Reconstructing cell histories in space with image-readable base editor recording. bioRxiv (Jan 2024). 10.1101/2024.01.03.573434, http://dx.doi.org/10.1101/2024.01.03.573434

4. Chan, M.M., Smith, Z.D., Grosswendt, S., Kretzmer, H., Norman, T.M., Adamson, B., Jost, M., Quinn, J.J., Yang, D., Jones, M.G., Khodaverdian, A., Yosef, N., Meissner, A., Weissman, J.S.: Molecular recording of mammalian embryogenesis. Nature 570(7759), 77–82 (Jun 2019). 10.1038/s41586-019-1184-5, https://doi.org/10.1038/s41586-019-1184-5

5. Chen, Z., Zhang, B., Tang, L., Gong, F., Wan, L., Ma, L.: Bayesian inference of lineage trees by joint analysis of single-cell multimodal lineage-tracing data with bilint (Sep 2025). 10.1101/2025.09.21.677669, http://dx.doi.org/10.1101/2025.09.21.677669

6. Evans, S.N., Winter, A.: Subtree prune and regraft: A reversible real tree-valued markov process. The Annals of Probability 34(3) (May 2006). 10.1214/009117906000000034, http://dx.doi.org/10.1214/009117906000000034

7. Felsenstein, J.: Evolutionary trees from dna sequences: A maximum likelihood approach. Journal of Molecular Evolution 17(6), 368–376 (Nov 1981). 10.1007/bf01734359, http://dx.doi.org/10.1007/BF01734359

8. Fu, W.Y.: Accelerated high-dimensional global optimization: A particle swarm optimizer incorporating homogeneous learning and autophagy mechanisms. Information Sciences 648, 119573 (Nov 2023). 10.1016/j.ins.2023.119573, http://dx.doi.org/10.1016/j.ins.2023.119573

9. Gong, W., Granados, A.A., Hu, J., Jones, M.G., Raz, O., Salvador-Martínez, I., Zhang, H., Chow, K.H.K., Kwak, I.Y., Retkute, R., Prusokas, A., Prusokas, A., Khodaverdian, A., Zhang, R., Rao, S., Wang, R., Rennert, P., Saipradeep, V.G., Sivadasan, N., Rao, A., Joseph, T., Srinivasan, R., Peng, J., Han, L., Shang, X., Garry, D.J., Yu, T., Chung, V., Mason, M., Liu, Z., Guan, Y., Yosef, N., Shendure, J., Telford, M.J., Shapiro, E., Elowitz, M.B., Meyer, P.: Benchmarked approaches for reconstruction of in vitro cell lineages and in silico models of c. elegans and m. musculus developmental trees. Cell Systems 12(8), 810–826.e4 (Aug 2021). 10.1016/j.cels.2021.05.008, https://doi.org/10.1016/j.cels.2021.05.008

10. Heaps, S.E., Nye, T.M., Boys, R.J., Williams, T.A., Embley, T.M.: Bayesian modelling of compositional heterogeneity in molecular phylogenetics. Statistical Applications in Genetics and Molecular Biology 13(5), 589–609 (Aug 2014). 10.1515/sagmb-2013-0077, http://dx.doi.org/10.1515/sagmb-2013-0077

11. Jones, M.G., Khodaverdian, A., Quinn, J.J., Chan, M.M., Hussmann, J.A., Wang, R., Xu, C., Weissman, J.S., Yosef, N.: Inference of single-cell phylogenies from lineage tracing data using cassiopeia. Genome Biology 21(1) (Apr 2020). 10.1186/s13059-020-02000-8, https://doi.org/10.1186/s13059-020-02000-8

12. Jones, M.G., Sun, D., Min, K.H.J., Colgan, W.N., Wang, H., Török, T., Cardoso, E.C., Tian, L., Weir, J.A., Chen, V.Z., Koblan, L.W., Yost, K.E., Mathey-Andrews, N., D’Souza, E., Russell, A.J., Stickels, R.R., Balderrama, K.S., Rideout, W.M., Dai, M., Marrero, G., Kumar, V., Saqi, A., Herzberg, B., Izar, B., Chang, H.Y., Lee, J.H., Jacks, T., Chen, F., Weissman, J.S., Yosef, N., Yang, D.: Spatiotemporal lineage tracing reveals the dynamic spatial architecture of tumour growth and metastasis. BioRxiv (Oct 2024). 10.1101/2024.10.21.619529, http://dx.doi.org/10.1101/2024.10.21.619529

13. Koblan, L.W., Yost, K.E., Zheng, P., Colgan, W.N., Jones, M.G., Yang, D., Kumar, A., Sandhu, J., Schnell, A., Sun, D., Ergen, C., Saunders, R.A., Zhuang, X., Allen, W.E., Yosef, N., Weissman, J.S.: High-resolution spatial mapping of cell state and lineage dynamics in vivo with PEtracer. Science 390(6770), eadx3800 (Oct 2025)

14. Li, K.R., Yu, P.L., Zheng, Q.Q., Wang, X., Fang, X., Li, L.C., Xu, C.R.: Spatiotemporal and genetic cell lineage tracing of endodermal organogenesis at single-cell resolution. Cell 188(3), 796–813.e24 (Feb 2025). 10.1016/j.cell.2024.12.012, http://dx.doi.org/10.1016/j.cell.2024.12.012

15. Liu, C., Zhou, X., Li, Y., Hittinger, C.T., Pan, R., Huang, J., Chen, X.x., Rokas, A., Chen, Y., Shen, X.X.: The influence of the number of tree searches on maximum likelihood inference in phylogenomics. Systematic Biology 73(5), 807–822 (Jun 2024). 10.1093/sysbio/syae031, http://dx.doi.org/10.1093/sysbio/syae031

16. Money, D., Whelan, S.: Characterizing the phylogenetic tree-search problem. Systematic Biology 61(2), 228 (Mar 2012). 10.1093/sysbio/syr097, http://dx.doi.org/10.1093/sysbio/syr097

17. Nye, T.M., Lio, P., Gilks, W.R.: A novel algorithm and web-based tool for comparing two alternative phylogenetic trees. Bioinformatics 22(1), 117–119 (Oct 2005). 10.1093/bioinformatics/bti720, https://doi.org/10.1093/bioinformatics/bti720

18. Pan, X., Li, H., Putta, P., Zhang, X.: Linrace: cell division history reconstruction of single cells using paired lineage barcode and gene expression data. Nature Communications 14(1) (Dec 2023). 10.1038/s41467-023-44173-3, http://dx.doi.org/10.1038/s41467-023-44173-3

19. Purvis, A., Garland, T.: Polytomies in comparative analyses of continuous characters. Systematic Biology 42(4), 569–575 (Dec 1993). 10.1093/sysbio/42.4.569, http://dx.doi.org/10.1093/sysbio/42.4.569

20. Robinson, D., Foulds, L.: Comparison of phylogenetic trees. Mathematical Biosciences 53(1-2), 131–147 (Feb 1981). 10.1016/0025-5564(81)90043-2, https://doi.org/10.1016/0025-5564(81)90043-2

21. Saitou, N., Nei, N.: The neighbor-joining method: a new method for reconstructing phylogenetic trees. Molecular Biology and Evolution (Jul 1987). 10.1093/oxfordjournals.molbev.a040454, http://dx.doi.org/10. 1093/oxfordjournals.molbev.a040454

22. Sashittal, P., Schmidt, H., Chan, M., Raphael, B.J.: Startle: A star homoplasy approach for crispr-cas9 lineage tracing. Cell Systems 14(12), 1113–1121.e9 (Dec 2023). 10.1016/j.cels.2023.11.005, http://dx.doi.org/10.1016/j.cels.2023.11.005

23. Short, S., García-Tejera, R., Schumacher, L.J., Coutu, D.L.: Next generation lineage tracing and its applications to unravel development. npj Systems Biology and Applications 11(1) (Jun 2025). 10.1038/s41540-025-00542-w, http://dx.doi.org/10.1038/s41540-025-00542-w

24. Slowinski, J.B.: Molecular polytomies. Molecular Phylogenetics and Evolution 19(1), 114–120 (Apr 2001). 10.1006/mpev.2000.0897, http://dx.doi.org/10.1006/mpev.2000.0897

25. Spanjaard, B., Hu, B., Mitic, N., Olivares-Chauvet, P., Janjuha, S., Ninov, N., Junker, J.P.: Simultaneous lineage tracing and cell-type identification using crispr–cas9-induced genetic scars. Nature Biotechnology 36(5), 469–473 (May 2018). 10.1038/nbt.4124, https://doi.org/10.1038/nbt.4124

26. Togkousidis, A., Kozlov, O.M., Haag, J., Höhler, D., Stamatakis, A.: Adaptive raxml-ng: Accelerating phylogenetic inference under maximum likelihood using dataset difficulty. Molecular Biology and Evolution 40(10) (Oct 2023). 10.1093/molbev/msad227, http://dx.doi.org/10.1093/molbev/msad227

27. Torres-Montúfar, A., Borsch, T., Ochoterena, H.: When homoplasy is not homoplasy: Dissecting trait evolution by contrasting composite and reductive coding. Systematic Biology 67(3), 543–551 (Jul 2017). 10.1093/sysbio/syx053, http://dx.doi.org/10.1093/sysbio/syx053

28. VanHorn, S., Morris, S.A.: Next-generation lineage tracing and fate mapping to interrogate development. Developmental Cell 56(1), 7–21 (Jan 2021). 10.1016/j.devcel.2020.10.021, http://dx.doi.org/10.1016/j.devcel.2020.10.021

29. Wagner, D.E., Klein, A.M.: Lineage tracing meets single-cell omics: opportunities and challenges. Nature Reviews Genetics 21(7), 410–427 (Jul 2020). 10.1038/s41576-020-0223-2, https://doi.org/10.1038/s41576-020-0223-2

30. Yang, J., Hou, L., Wang, X., Zhang, N., Bian, Y., Lu, Z., Chen, Y., Xie, D., Fang, Y., Wang, K., Wan, R., Jin, Y., Chen, Y., Cai, X., On Lee, L.T., Hu, Z., Ji, H.: Spatiotemporal lineage mapping of tumor immune escape with etracer (Aug 2025). 10.1101/2025.08.06.668639, http://dx.doi.org/10.1101/2025.08.06.668639

31. Zafar, H., Lin, C., Bar-Joseph, Z.: Single-cell lineage tracing by integrating CRISPR-cas9 mutations with transcriptomic data. Nature Communications 11(1) (Jun 2020). 10.1038/s41467-020-16821-5, https://doi.org/10.1038/s41467-020-16821-5

32. Zou, Y., Zhang, Z., Zeng, Y., Hu, H., Hao, Y., Huang, S., Li, B.: Common methods for phylogenetic tree construction and their implementation in r. Bioengineering 11(5), 480 (May 2024). 10.3390/bioengineering11050480, http://dx.doi.org/10.3390/bioengineering11050480

