## Supplementary Information for "Integrative Inference of Spatially Resolved Cell Lineage Trees using LineageMap"

**Abstract.** Understanding the spatio-temporal processes of tissue growth, including how new cell types emerge and how cells form the tissue architecture, is a fundamental problem in biology. The emerging spatially resolved lineage tracing data, where three modalities, which are lineage barcodes, gene expression profiles and spatial locations are measured for each single cell, provides unprecedented opportunity to understand these processes. Computational methods that take advantage of all three modalities to reconstruct cell lineage tree and ancestral cell states and locations are needed. We introduce LineageMap, a hybrid lineage inference algorithm that integrates the scalability of distance-based tree reconstruction methods with the Interpretability of likelihood-based methods under a unified probabilistic framework. The input to LineageMap is spatially resolved lineage tracing data, where for each single cell, the gene expression, lineage barcode and spatial locations are available. LineageMap enables accurate, interpretable, and scalable inference of high-resolution lineage trees as well as locations of ancestral cells from multi-omic single-cell data. Across simulated and experimental datasets, LineageMap consistently outperforms existing methods in reconstructing both global topology and local clonal structure, while revealing biologically coherent spatiotemporal trajectories. Our framework bridges molecular lineage tracing with spatial and transcriptomic information, advancing computational reconstruction of dynamic cellular ancestries at the tissue scale. LineageMap is available at: https://github.com/ZhangLabGT/LineageMap.

**Keywords:** Cell lineage inference · Spatial transcriptomics · Evolutionary modeling · Maximum likelihood.

### 1 Supplementary Figures

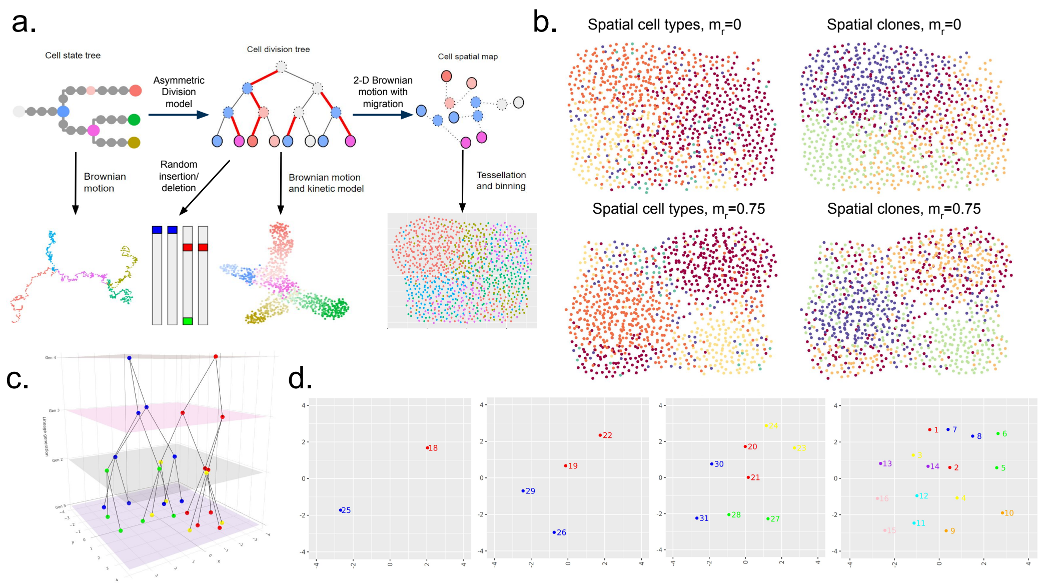


Supplementary Figure 1: (a) SpaTedSim simulation framework. SpaTedSim takes the input of a cell state tree, a cell and a cell division tree, and outputs paired lineage barcodes, gene expressions and spatial coordinates. (b) Visualization of SpaTedSim generated datasets, colored by cell type or clone ID. The top row shows a low migration rate, better clone clusters, and worse cell-type clusters; the bottom row shows a high migration rate, worse clone clusters, and better cell-type clusters. (c) A 3D visualization of the simulated lineage tree projected into spatial and temporal dimensions. X and Y axes represent the spatial coordinates of the cells, while Z axis represents the No. generation from the root cell, and cells are colored with their clonal id. (d) The 2D visualization of each generation from generation 2 to generation 5. The nodes colored based on their clonal relationship and are numbered following the post order of the tree in (c).

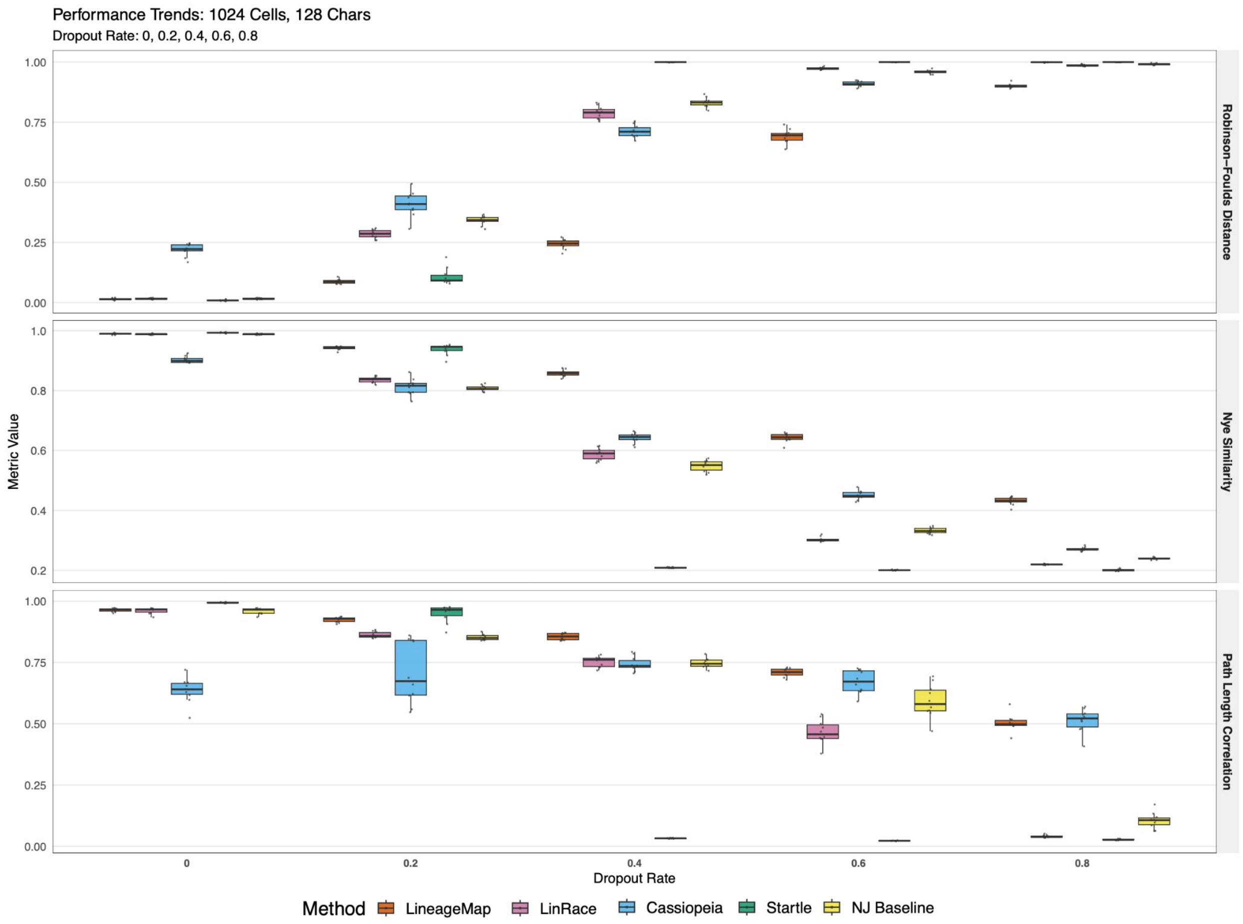


Supplementary Figure 2: Experiments to evaluate the robustness of tools across multiple dropout rate setting in simulated data. For RF distance, lower value means better performance. For the Nye similarity and path length correlation, higher values mean better performance. All tools' performance decreased as dropout rate increasing, while LineageMap remains to be relatively stable and have better performance, and especially outperformance other tools in the situation of high dropout rate datasets.

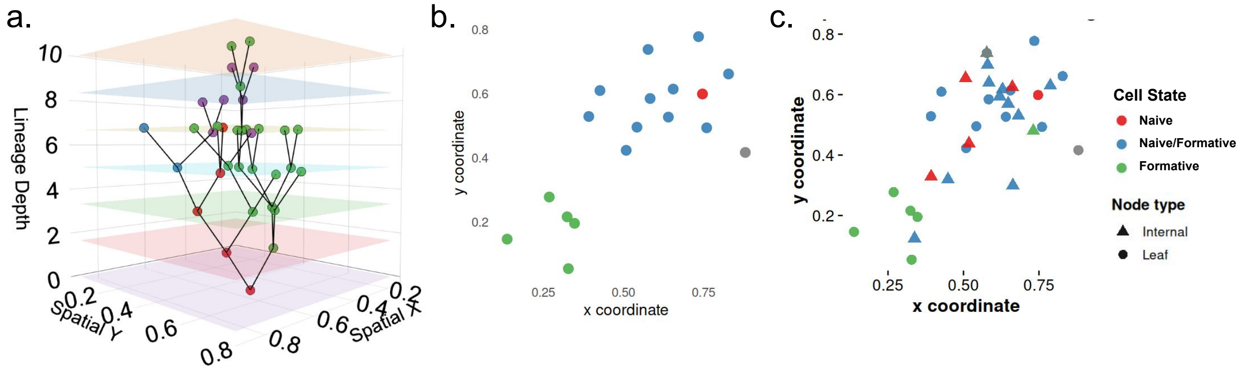


Supplementary Figure 3: LineageMap infers spatially-resolved lineage on baseMEMOIR mESC data (colony 4). (a) A ploty 3D visualization of the reconstructed lineage tree projected into spatial and temporal dimensions. X and Y axes represent the spatial coordinates of the cells, while Z axis represents the evolutionary depth from the root, and cells are colored with cell types. (b) 2D visualization of spatial coordinates of observed leaf cells, colored by cell states. (c) 2D visualization of spatial coordinates of both observed leaf cells, and inferred ancestral cells, colored by cell states. The cell states and spatial coordinates of ancestral cells are inferred using LineageMap's likelihood function.

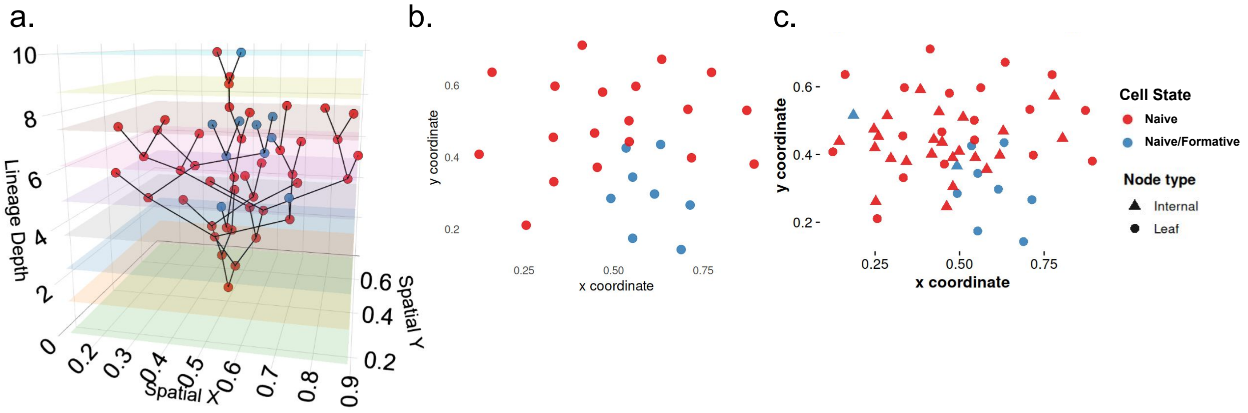


Supplementary Figure 4: LineageMap infers spatially-resolved lineage on baseMEMOIR mESC data (colony 5). (a) A ploty 3D visualization of the reconstructed lineage tree projected into spatial and temporal dimensions. X and Y axes represent the spatial coordinates of the cells, while Z axis represents the evolutionary depth from the root, and cells are colored with cell types. (b) 2D visualization of spatial coordinates of observed leaf cells, colored by cell states. (c) 2D visualization of spatial coordinates of both observed leaf cells, and inferred ancestral cells, colored by cell states. The cell states and spatial coordinates of ancestral cells are inferred using LineageMap's likelihood function.

#### 2 Derivation of tri-modality likelihood function

##### 2.1 Barcode likelihood

For locus $l$, edits occur irreversibly from the unedited state. Let $\lambda_{lk}$ be the rate of editing from un-mutated state 0 to a mutated state $k$, with total mutation rate $\lambda_l = \sum_{k=1}^{K_l} \lambda_{lk}$. The transition probability matrix between barcode states can be written as:

$$P_l(t) = \begin{bmatrix} e^{-\lambda_l t} & \frac{\lambda_{l1}}{\lambda_l}\left(1-e^{-\lambda_l t}\right) & \cdots & \frac{\lambda_{lK_l}}{\lambda_l}\left(1-e^{-\lambda_l t}\right) \\ 0 & 1 & \cdots & 0 \\ \vdots & \vdots & \ddots & \vdots \\ 0 & 0 & \cdots & 1 \end{bmatrix}.$$

##### 2.2 Modeling cell migration into Brownian motion model

Here we give detailed derivations to calculating Gaussian message passing using OU model. We observe spatial coordinates $\{\mathbf{s}_i\}_{i \in d}$ at the leaves $L$ of a rooted tree $T = (V, E)$. We assume an edgewise, conditional Gaussian model

$$\mathbf{s}_v \mid \mathbf{s}_u \ \sim \ \mathcal{N}\big(\mathbf{m}(\mathbf{s}_u, t_{uv}),\ \Sigma_s(t_{uv})\big) \qquad \text{for } (u \to v) \in E,$$

and a root prior $\mathbf{s}_r \sim \mathcal{N}(\mu_0, \Sigma_0)$. To make sure that the likelihood is tractable by the Gaussian message passing algorithm, we use the Ornstein–Uhlenbeck process to model cell migration towards specific tissue regions.

**Affine transitions preserve Gaussianity**

If for every edge $(u \to v)$ the transition has the *affine linear* form

$$\mathbf{m}(\mathbf{s}_u, t) = A(t)\,\mathbf{s}_u + \mathbf{b}(t)\ , \qquad \Sigma_s(t) = S_e(t)\ ,$$

with $A(t) \in \mathbb{R}^{d \times d}$, $\mathbf{b}(t) \in \mathbb{R}^d$, and $S_e(t)$ positive definite, then

$$\mathbf{s}_v \mid \mathbf{s}_u \sim \mathcal{N}\big(A\mathbf{s}_u + \mathbf{b},\ S_e\big)$$

and the family of Gaussian subtree messages is *closed* under the upward convolution

$$m_{c \to u}(\mathbf{s}_u) \ = \ \int p(\mathbf{s}_v \mid \mathbf{s}_u)\, p(\text{data in subtree } c \mid \mathbf{s}_v)\, d\mathbf{s}_v,$$

so message passing remains in the Gaussian family and is therefore tractable. The multivariate OU process $\mathbf{s}(t) \in \mathbb{R}^d$ is defined by the SDE

$$d\mathbf{s}(t) = -K\big(\mathbf{s}(t) - \mu\big)\, dt + L\, d\mathbf{W}(t),$$

where $K \in \mathbb{R}^{d \times d}$ is a positive-definite rate matrix (pulling toward the attractor $\mu \in \mathbb{R}^d$), $L \in \mathbb{R}^{d \times r}$ is a dispersion matrix, and $\Sigma := LL^\top$ is the diffusion matrix. For an edge of length $t$ the transition distribution is Gaussian:

$$\mathbf{s}_v \mid \mathbf{s}_u \ \sim \ \mathcal{N}\big(A(t)\,\mathbf{s}_u + \mathbf{b}(t),\ S_e(t)\big),$$

with

$$A(t) = e^{-Kt}, \qquad \mathbf{b}(t) = \big(I - e^{-Kt}\big)\mu,$$

and

$$S_e(t) \ = \ \int_0^t e^{-Ks}\, \Sigma\, e^{-K^\top s}\, ds.$$

Thus the OU transition is affine in the parent state and the transition covariance is the above integral. We will formally define the state-dependent OU model and the computation of marginal likelihood on a lineage tree.

#### 3 Dynamic threshold selection for constructing the barcode similarity graph

To identify coarse sublineages prior to local tree refinement, we apply Louvain community detection to a similarity graph derived from lineage barcode comparisons. A key parameter in this procedure is the similarity threshold used to determine whether an edge is included in the graph. Because lineage barcodes obtained from CRISPR / MEMOIR-like systems can vary substantially in mutation sparsity and dropout frequency, a fixed threshold is often suboptimal. We therefore developed a dynamic, data-driven procedure that adapts the threshold to the observed distribution of barcode similarities.

##### Dropout Rate Estimation

Let $X \in \mathbb{R}^{n\times m}$ denote the barcode matrix for $n$ cells and $m$ target sites, where each entry may encode a mutated state, an unmutated state, or a dropout indicator (e.g. "–"). We define the global dropout rate as

$$\delta \ = \ \frac{1}{nm}\big|\{\, X_{ij} = \text{dropout}\,\}\big|,$$

representing the fraction of missing or unreadable barcode sites. High dropout reduces the effective resolution of pairwise barcode comparisons and lowers similarity values, motivating adaptive thresholding.

##### Similarity Matrix Construction

We compute a dropout-aware pairwise distance matrix $D$ using a custom function `DropoutDistMatrix`, and convert distances into similarities via

$$S_{ij} \ = \ 1 - D_{ij}.$$

Let $\mathcal{S} = \{S_{ij} \mid i < j\}$ denote the multiset of pairwise similarities. In practice, $\mathcal{S}$ often exhibits a bimodal structure, with one mode corresponding to unrelated cells and another corresponding to sublineages sharing common ancestral mutations.

##### Otsu Thresholding for Bimodal Similarity Distributions

When $\mathcal{S}$ is bimodal, we apply Otsu thresholding to automatically determine a similarity cutoff $\tau$. Otsu's method selects the value $\hat{s}$ that maximizes the between-class variance of a two-class partition of $\mathcal{S}$:

$$\hat{s} \ = \ \arg\max_{s}\Big[\, w_B(s)\, w_F(s)\, \big(\mu_B(s) - \mu_F(s)\big)^2 \Big],$$

where $w_B(s)$ and $w_F(s)$ are the proportions of similarities below and above $s$, respectively, and $\mu_B(s)$ and $\mu_F(s)$ are their means. The resulting similarity threshold is set to

$$\tau \ = \ \hat{s}.$$

Otsu's method robustly separates similarities arising from noise and dropout from similarities reflecting true shared ancestry, without requiring user-specified parameters.

##### Quantile-Based Thresholding for High-Dropout Regimes

In datasets with extreme dropout, the similarity distribution may collapse into a unimodal or heavy-tailed form, making Otsu thresholding unstable. In these cases we use a quantile-based cutoff

$$\tau \ = \ \text{Quantile}\big(\mathcal{S},\, q(\delta)\big),$$

where the quantile level is adjusted according to dropout:

$$q(\delta) \ = \ 0.90 + 0.05\,\delta.$$

This scheme becomes more conservative (i.e. increases the required similarity) as dropout rises, preventing spurious edges caused by missing data.

##### Dropout-Adjusted Analytic Threshold (Fallback Rule)

As an additional safeguard, if neither Otsu nor quantile thresholding yields a stable cutoff (e.g. for extremely small cell counts), we employ a simple analytic rule derived from empirical behavior across simulated and real datasets:

$$\tau \ = \ 0.12 + 0.4\,\delta.$$

This fallback rule preserves coarse sublineage separation under degenerate conditions.

##### Graph Construction and Louvain Clustering

Given the dynamically chosen threshold $\tau$, we construct an undirected weighted graph $G$ whose vertices correspond to cells and whose edges connect all pairs $(i, j)$ with $S_{ij} > \tau$. Edge weights are set to the underlying similarity values $S_{ij}$. We then apply Louvain community detection with resolution parameter $\gamma = 1$ to obtain initial sublineage groups. These groups form the backbone structure that is subsequently refined by the local search procedure described in the main text.

### 4 Simulating paired gene expression, lineage barcodes, and spatial coordinates using SpaTedSim

To characterize the spatio-temporal dynamics of cells' transcriptomes, and provide a tool to test computational methods for multimodal data integration, we developed SpaTedSim, a spatio-temporal dynamics simulation of single cells. SpaTedSim generates paired single-cell RNA counts, spatial coordinates, and lineage barcodes simultaneously, by simulating realistic biological events including cell symmetric and asymmetric divisions, and cell migrations. SpaTedSim can tune the parameters for cell division and migration events that lead to changes in the spatio-temporal correlation of cells' gene expressions, accommodating various biological circumstances. SpaTedSim can test different types of computational tasks, including trajectory inference, lineage inference, spatial transcriptome mapping, etc. SpaTedSim can also fit real ST data and generate synthetic datasets that share realistic spatial gene expression with clonal information.

#### 4.1 Simulating cells' lineage barcode and gene expressions on the cell division tree

we adopt the TedSim framework: We start with two tree structures: a cell state tree, indicating the developmental trajectory of cell types; and a cell division tree, indicating the cell division history. TedSim first runs a Brownian motion model on the cell state tree to obtain the mean latent space representations for each cell state. TedSim then runs another Brownian motion model on the cell division tree to obtain the latent space representations and then transform the latent space into gene expression levels for each cell. A cell's latent space representation can be regarded as a weighted sum of the state mean representations and the cell's lineage representations. We adopt the asymmetric division model to determine the cell states of all cells while performing Brownian motion on the lineage tree. Starting from the root of the cell lineage tree, a cell can divide either symmetrically into two cells of the same state as their parent or asymmetrically, which means one child cell stays at the same state and the other child cell shifts to a future state on the cell state tree. The leaf cells on the cell division tree are considered present-day cells, and the gene expression profiles of the leaf cells are the output transcriptomic data for a scRNA-seq experiment. A detailed description of the simulation of gene expression and lineage barcode can be found in TedSim [1].

#### 4.2 Simulating spatial coordinates using SpaTedSim

Given a lineage tree with $n_{\text{cells}}$ tips and $n_{\text{gen}}$ generations, SpaTedSim initializes the root cell at the spatial origin and iteratively expands the population through simulated cell divisions. At each generation, the spatial coordinates of newly produced cells are determined by three key components:

a) **Cell division:** Each parent cell generates two children whose positions are drawn from an angular displacement model with distance decreasing across generations. If $(x_p, y_p)$ is the parent coordinate, the children positions are

$$(x_p \pm r\cos\theta + \epsilon_x,\ y_p \pm r\sin\theta + \epsilon_y),$$

where $\theta \sim U(0, 2\pi)$, $r = \frac{\text{division\_radius}}{\sqrt{4^{1.5(g-1)}}}$ decays with generation $g$, and the noise terms follow $\epsilon_x, \epsilon_y \sim \mathcal{N}(0, \sigma^2)$.

b) **Cell migration:** Cells may undergo active migration with probability proportional to a generation-dependent rate. Migration is guided by a kernel density estimate (KDE) of (i) same-type cells and (ii) background cells. Candidate migration destinations are sampled from a local grid around each cell and evaluated using a density contrast function:

$$\Delta(x, y) = \rho_{\text{type}}(x, y) - \rho_{\text{background}}(x, y).$$

Cells preferentially relocate to regions maximizing this contrast, with additional stochastic noise.

c) **Boundary enforcement:** To prevent cells from leaving the defined spatial domain $[x_{\min}, x_{\max}] \times [y_{\min}, y_{\max}]$, boundary-violating proposals are projected back into feasible space. Projection uses a KDE-weighted sampling scheme within a local circular region centered at the boundary location.

**Density-Guided Gradient Drift**

To avoid degenerate clustering and encourage spatially realistic tissue distributions, SpaTedSim implements a low-density drift process. For each cell, local neighborhoods are examined using a fine grid; the cell then moves toward regions of lower density with probability proportional to the inverse KDE density:

$$p(x, y) \propto \max(\rho) - \rho(x, y).$$

Gaussian noise is applied after each drift step.

**Post-Processing: Distance Constraints**

SpaTedSim applies two geometric post-processing steps after early generations:

- **Minimum-distance repulsion:** Ensures that no two cells lie closer than a user-specified threshold. Iteratively, pairs within the threshold repel each other using normalized displacement vectors.
- **Maximum-distance attraction:** Ensures that a cell's $k$-th nearest neighbor is not excessively far away, promoting spatial continuity.

Detailed Pseudocode of the post processing steps can be found below.

##### 4.3 Post-processing of simulated spatial coordinates of cells

Post-processing has two steps to make the overall spatial distribution of cells more uniform: 1. Pull the outlier cells that are too far away from the neighbors; 2. Push the cells too close to each other based on a given minimum distance. Here are the pseudocodes that describes the two steps:

**Pseudocode for pulling cells based on maximum distance constraint**

- **Input:**
  - spatial coordinates of cells (x, y)
  - *max_distance*: Maximum allowed distance between points
  - $k$: The $k$-th closest point to check (default is 3)
  - *iterations*: Number of iterations to perform (default is 100)

- *step_size*: Step size for adjusting points (default is 0.05)

– **Procedure:**

a) For each iteration from 1 to *iterations*:

(a) Recalculate the distance matrix $D$ between all cells $D = \{D_{ij} = ||(x_i - x_j)^2 + (y_i - y_j)^2||_2\}$

(b) For each cell $(x_j, y_j)$:

i. Find the $k$-th closest cell to cell $j$ in $D_{*,j}$

ii. If the distance to the $k$-th closest cell exceeds *max_distance*:

A. Identify the index $l$ of the $k$-th closest cell

B. Compute the vector $\vec{v}$ from cell $j$ to cell $l$

C. Normalize $\vec{v}$ and scale it by *step_size*

D. Adjust cell $j$'s spatial coordinates by subtracting the components of the scaled vector: $(x'_j, y'_j) = (x_j, y_j) - \mathit{step\_size} \cdot \frac{\vec{v}}{|\vec{v}|}$

(c) Recalculate the distance matrix after the adjustment

b) If the distance of all points to their $k$-th closest neighbor is less than or equal to *max_distance*, break the loop and return the updated spatial coordinates of cells

– **Output:**

- Adjusted spatial coordinates of cells $(x', y')$.

**Pseudocode for pushing cells based on minimum distance constraint**

– **Input:**

- Spatial coordinates of cells (x, y)
- *min_distance*: Minimum allowed distance between points
- *iterations*: Maximum number of iterations to perform (default is 20)
- *step_size*: Step size for adjusting points (default is 0.05)

– **Procedure:**

a) For each iteration from 1 to *iterations*:

(a) Recompute the distance matrix between the spatial coordinates of all cells: $D = \{D_{ij} = ||(x_i - x_j)^2 + (y_i - y_j)^2||_2\}$

(b) For each cell $(x_j, y_j)$:

i. Identify all cells closer to $j$ than *min_distance* but not at a zero distance

ii. For each point $k$ that is too close:

A. Compute the vector $\vec{v}$ from point $j$ to point $k$

B. Normalize $\vec{v}$ and scale it by *step_size*

C. Adjust spatial coordinates of cell $j$ by adding the components of the scaled vector $(x'_j, y'_j) = (x_j, y_j) + \mathit{step\_size} \cdot \frac{\vec{v}}{|\vec{v}|}$

D. Adjust spatial coordinates of cell $k$ by subtracting the components of the scaled vector: $(x'_k, y'_k) = (x_k, y_k) - \mathit{step\_size} \cdot \frac{\vec{v}}{|\vec{v}|}$

(c) Recompute the distance matrix after adjustments

b) If all pairwise distances are greater than or equal to *min_distance*, break the loop and return the updated spatial coordinates of cells

– **Output:**

- Adjusted spatial coordinates of cells $(x', y')$.

### 5 Experimental Details

#### 5.1 SpaTedSim simulation procedure and settings

(1) Varying variables in the simulation include:
Migration rate: $\mu_m$ = [0,1].
Dropout: $p_d$ = [0,0.2,0.4,0.6,0.8].

Number of cells: 128 or 1024.
(2) Simulation parameters for simulating gene expressions, lineage barcodes and spatial coordinates using SpaTedSim:

Table 1: Simulation settings for gene expression in SpaTedSim

| | |
|---|---|
| Number of genes (ngenes) | 500 |
| Number of Identity Vectors ($N_{IV}$) | 30 |
| State Identity Vector stepsize (*step*) | 0.5 |
| Maximum number of state shifts for one division ($max\_walk$) | 6 |
| Asymmetric division rate (p_a) | 0.6 |
| Identity Vector center (starting value for diff-IF) | 1 |
| Number of diff-Identity Vectors ($N_{diff}$) | 20 |
| nondiff-SIV standard deviation ($\sigma$) | 0.5 |
| Probability of nonzero gene effect (ge_prob) | 0.3 |
| Probability of outlier gene (prob_hge) | 0.03 |
| Mean of capture efficiency $\alpha$ (alpha_mean) | 0.1 |
| Standard deviation of capture efficiency $\alpha$ (alpha_sd) | 0.02 |

Table 2: Simulation settings for lineage barcodes in SpaTedSim

| | |
|---|---|
| Number of characters | 16,128 |
| Excision dropout rate | 0 |
| Distribution for mutation sampling | Exponential |

Table 3: Simulation settings for spatial coordinates in SpaTedSim

| | |
|---|---|
| Standard deviation of 2-D Brownian motion($\sigma_2$) | 0.6 |
| Division radius ($r_d$) | 3(128 cells), 6(1024 cells) |
| Migration radius ($r_m$) | 6(128 cells), 12(1024 cells) |
| Spatial coordinate range (both x and y) | (-3,3) for 128 cells, (-6,6) for 1024 cells |
| min_distance | 0.2 |
| max_distance | 1 |

#### 5.2 Detailed settings of running LineageMap on SpaTedSim datasets

To evaluate the performance of our method under controlled conditions with known ground truth, we applied LineageMap to synthetic lineage datasets generated using SpaTedSim. SpaTedSim produces spatially resolved lineage barcodes, simulated cell states along predefined developmental lineages, and realistic spatial arrangements of cells following an Ornstein–Uhlenbeck diffusion model. The resulting data provide an ideal benchmark for assessing both the topological and spatial accuracy of inferred lineage trees. To systematically evaluate lineage reconstruction performance across different levels of barcode complexity, sequencing sparsity, and dataset size, we generated a grid of SpaTedSim datasets varying three key parameters:

– **Number of cells:** $N \in \{128, 1024\}$
– **Number of barcode sites:** $K \in \{16, 128\}$
– **Character dropout rate:** $d \in \{0.0, 0.2, 0.4\}$

For each $(N, K, d)$ combination, we generated **10 independent replicates**.
This design allows controlled evaluation of:

– the effect of dataset size (128 vs. 1024 cells),
– the effect of barcode richness (16 vs. 128 sites),
– robustness to barcode dropout (0–40%),
– variability across replicate simulations.

All datasets contain ground-truth lineage trees, barcode matrices, spatial coordinates, and cell-state labels generated by SpaTedSim. All simulated datasets will be randomized by row for avoiding running bias caused by organized order. For different numbers of cells and different numbers of target sites, we also tested the running time efficiency of LineageMap. Given $(N, K)$, the time complexity of LineageMap is $O(N^2K)$, and below is the actual running time on different settings:

Table 4: Average runtime (seconds) across datasets of varying input size.

| Number of Cells ($N$) | Number of target sites($K$) | Runtime (s) |
|---|---|---|
| 128 | 16 | 5.15 |
| 512 | 16 | 79.20 |
| 1024 | 16 | 315.56 |
| 2048 | 16 | 1283.02 |
| 4096 | 16 | 5310.16 |
| 1024 | 16 | 315.56 |
| 1024 | 32 | 565.98 |
| 1024 | 64 | 1052.55 |

##### 5.3 Detailed settings of running LineageMap on baseMEMOIR mESC data

To evaluate lineage reconstruction performance on the baseMEMOIR mouse embryonic stem cell (mESC) dataset, we ran LineageMap on selected colonies (IDs 2, 4, 5) following the procedure below.

###### Data Preparation

a) **Lineage Trees:** Ground-truth lineage trees for each colony were obtained from BEAST analysis in Newick format.
b) **Spatial Coordinates:** Cell location data (x, y) and cell states were read from CSV files and filtered to include only sequenced cells. Cell IDs were sorted by lineage node for consistency.
c) **Barcode Sequences:** Single-cell barcodes were read in Nexus format. Unknown barcode entries ("?") were replaced with "-" for compatibility with LineageMap.
d) **State Lineages:** Three sets of developmental states were defined for inference:

$$\text{state\_lineages} = \{[1, 4, 5], [1, 2, 3], [2, 3, 4, 5]\}.$$

**Running LineageMap**

LineageMap was run with the following configurations:

a) **Backbone Tree Construction:**
   - Backbone tree built using with neighbor-joining median method (`backbone_type = "NJ_median"`).
   - Parallelization: `outer_cores = 12`, `inner_cores = 5`.
   - hyperparameters of the likelihood function: $\lambda_1 = 0.05$, $\lambda_2 = 0.1$, $\alpha = 1$.
   - Threshold for building the barcode similarity graph: 0.2.

b) **Post-Processing:**
   - Reroot the tree to balance left and right clones and converted inferred trees to ultrametric form.

##### 5.4 Details of running state-of-the-art methods

To evaluate the performance of inferring lineage tree, we compared LineageMap with the neighbor-joining baseline method and other state-of-the-art methods, including LinRace, Cassiopeia and Startle.

All inputs will be preprocessed to meet the requirement of these tools. For LinRace, we set all parameters to be default. For Cassiopeia, we tested its basic Vanilla mode and other hyperparameters are set to be default. For Startle, we set LARGE parsimony problem mode, and used classic neighbor-joining method as its initialization seed tree. Other hyperparameter of Startle are in default.

##### 5.5 Evaluation metrics for benchmarking lineage tree inference methods

To quantitatively assess how well an inferred lineage tree recapitulates the ground-truth structure, we evaluated each reconstructed tree using three complementary metrics: the Robinson–Foulds (RF) distance, the Nye similarity index, and the cophenetic path correlation. These metrics jointly characterize topological accuracy and preservation of pairwise cell relationships.

a) **Robinson–Foulds (RF) Distance** The Robinson–Foulds distance measures the topological disagreement between two trees by counting the number of bipartitions (splits) present in one tree but absent in the other. We report the *normalized* RF distance so that values lie in the interval $[0, 1]$, where 0 indicates identical topologies and 1 indicates maximal disagreement.
   Given a ground-truth tree $T_{\text{gt}}$ and an inferred tree $T$, the metric is computed as:

$$\text{RF}(T_{\text{gt}}, T) = \frac{\#\{\text{splits present in exactly one of } T_{\text{gt}}, T\}}{\#\{\text{splits in } T_{\text{gt}} \cup T\}}.$$

b) **Nye Similarity**
   The Nye similarity index provides a complementary measure of topological agreement by quantifying the proportion of internal splits shared between two trees. In contrast to the RF distance, which counts disagreements, the Nye similarity increases as the two trees share more structure. Values lie in $[0, 1]$, with 1 indicating identical branching topology.
   Reporting both RF distance and Nye similarity allows simultaneous evaluation of mismatches and shared structure.

c) **Cophenetic Path Correlation**
   While RF and Nye metrics evaluate discrete topological features, they do not assess the preservation of continuous lineage distances. To evaluate consistency of pairwise cell relationships, we compute the Pearson correlation between the cophenetic distance matrices of the inferred and ground-truth trees.
   Let $C_{\text{gt}}$ and $C$ denote the cophenetic matrices of $T_{\text{gt}}$ and $T$ respectively. After restricting to shared tip labels, we compute:

$$\text{PathCorr}(T_{\text{gt}}, T) = \text{cor}\left(\text{vec}(C_{\text{gt}}),\ \text{vec}(C)\right),$$

   where $\text{vec}(\cdot)$ denotes vectorization of the upper-triangular entries.
   This metric quantifies how well the inferred tree preserves the relative lineage distances among all pairs of cells.

#### References

1. Pan, X., Li, H., Zhang, X.: TedSim: temporal dynamics simulation of single-cell RNA sequencing data and cell division history. Nucleic Acids Research **50**(8), 4272–4288 (Apr 2022). https://doi.org/10.1093/nar/gkac235, `https://doi.org/10.1093/nar/gkac235`
